# Impact of conscious awareness on pupillary response to faces

**DOI:** 10.1101/2020.11.11.377614

**Authors:** Yung-Hao Yang, Hsin-I Liao, Shigeto Furukawa

## Abstract

Pupillary response reflects not only ambient light changes but also top-down factors. Nevertheless, it remains inconclusive whether the conscious awareness modulates the pupillary response. We investigated pupillary responses to faces under different conscious conditions using continuous flash suppression (CFS). In Experiment 1 and 2, we used a breaking-CFS procedure in which participants had to detect the face from suppression. Results showed that the pupil constricted more to upright faces than to inverted faces before the face was detected, suggesting that pupillary responses reflect face processing entering consciousness. In Experiment 3 and 4, we used a fixed duration-CFS procedure with both objective performance and subjective reports. Different pupillary responses were observed only when the participant was aware of the face. These findings imply that the conscious awareness is critical for modulating autonomic neural circuits of the pupillary function. The corresponding pupillary responses may reflect dynamic processes underlying conscious awareness.

## Introduction

The pupils primary function is to adjust the intensity of light inputs to the retina by changing its size in response to ambient light variations, which is referred to as the pupillary light response (de Groot & Gebhard, 1952). In addition to luminance changes, pupils also respond to other afferent visual features such as spatial gratings, motion, and chromatic stimuli under constant luminance situations (Barbur, Harlow, & Sahraie, 1992; Gamlin, Zhang, Harlow, & Barbur, 1998; Sahraie & Barbur, 1997). For complex visual information, a recent study reported that upright faces trigger stronger pupillary constriction than upside-down ones. This face inversion effect is species-specific for human faces in human observers (Conway, Jones, DeBruine, Little, & Sahraie, 2008). This suggests that complex objects such as faces could be represented as a kind of afferent visual feature to induce a spontaneous pupillary response. Given the close links between pupillary response and visual attributes, studies have assumed that pupillary responses can be a noninvasive, indirect, and objective measurement for investigating both low-level and high-level visual processing (Binda & Murray, 2014; Mathôt & Van der Stigchel, 2015).

The neural circuit of the pupillary response to visual features is underlain by projection from the extrastriate visual cortices to the Edinger–Westphal (EW) nucleus (Barbur, 2004). The EW nucleus contains preganglionic parasympathetic neurons, which project to postganglionic ciliary ganglion to induce pupil constriction (i.e., miosis) via innervating iris sphincter muscle. For the pupillary light response, when a transient luminance change occurs, the EW nucleus receives inputs from the olivary pretectal nucleus (OPN), which is activated by afferent signals from retinal ganglion cells responding to retinal photoreceptors. For the pupillary response to other visual features, according to Barbur’s model (2004), when there is no transient luminance change, the extrastriate visual cortices send a steady-state inhibitory signal to the EW nucleus to decrease the efferent signal to the iris sphincter muscle and thus maintain the pupil size in a normal range. When the processing of various visual features, such as color, spatial gratings, and motion, disturbs neural activities in the extrastriate visual cortices, the steady-state inhibition to the EW nucleus is decreased and thus the pupil constricts.

An issue in the above sophisticated neural model of the pupillary response to visual features is whether the pupillary response is modulated by top-down factors such as attention and/or consciousness, as the pupillary response is not merely a reflex. Indeed, it has been shown that the pupil constricts more when participants focus attention on bright surfaces (Binda, Pereverzeva, & Murray, 2014) or features (Mathôt, Dalmaijer, Grainger, & Stigchel, 2014) than on dark counterparts, suggesting that the pupillary light response is modulated by covert visual attention when the luminance input to the retina is fixed. On the other hand, it remains controversial whether conscious awareness modulates pupillary responses to visual features. In other words, is conscious awareness of the visual stimuli necessary for a pupillary response to them? Neurological studies suggest that conscious awareness formation relies on global recurrent neuronal processing from the prefrontal cortex (Dehaene, Changeux, Naccache, & Sergent, 2006; Del Cul, Dehaene, Reyes, Bravo, & Slachevsky, 2009; Lamme, 2006; but see Tong, 2003; Zeki, 2003). In this case, it is surmised that consciousness is not necessary for pupillary responses to visual features since the process that triggers them is completed within the visual cortices before conscious awareness is reached. Along this line, evidence shows that blindsight patients with V1 damage yet exhibit pupillary constriction in response to achromatic gratings and red-colored patches in their blind fields (Weiskrantz, Cowey, Le Mare, 1998; Weiskrantz, Cowey, Barbur, 1999). Moreover, by presenting facial expressions to the blind visual fields, an invisible fearful face induced more substantial pupillary dilation than a happy face in affective blindsight patients (Tamietto et al., 2009). These findings imply that pupillary responses reflect the processing of visual inputs, including colors and faces, in the absence of conscious awareness.

In contrast, studies using neurologically intact participants have shown that the pupil reflects the conscious percept when two competing visual inputs are dichoptically projected to the two eyes under binocular rivalry. In binocular rivalry, the conscious representation of a stimulus usually fluctuates across time, given the retina inputs are constant. Findings show that pupillary response corresponds to the dominant percept in binocular rivalry, in which the pupil constricts when the brighter percept is dominant and dilates when the darker percept is (Fahle, Stemmler, Spang 2011; Naber, Frassle, & Einhäuser, 2011; Schütz, Busch, Gorka, Einhäuser, 2018). Furthermore, pupillary light responses are inhibited when observers are unaware of luminance changes, although the physical luminance input to the retina indeed changes during the suppression phase (Barany & Hallden, 1948; Lorber, Zuber, & Stark, 1965). However, since the stimuli reciprocally compete with each other in the binocular rivalry condition, it is hard to determine whether the effect is “due to the effectiveness of the suppressed stimulus or the ineffectiveness of the suppressing stimulus” (Jiang, Costello, and He, 2007). The findings that pupillary responses reflecting the dominant conscious percept under this specific condition do not necessarily imply that consciousness is necessary for pupillary responses to visual features.

Recently, Sperandio, Bond, and Binda (2018) adopted the continuous flash suppression paradigm (CFS; Fang & He, 2005; Tsuchiya & Koch, 2005), a more sustainably interocular suppression method, to test whether a conscious appraisal is necessary for pupillary responses to scene perception related to brightness. In their study (Sperandio et al., 2018), a picture of the sun or a scrambled counterpart was presented to one eye, and a series of high-contrast masks (CFS condition) or a blank (non-CFS condition) was presented to the other eye to render the tested picture invisible or visible, respectively. Since the pictures of the sun and scrambled counterpart are both suppressed by the similar masks (in the CFS condition), unlike conventional binocular rivalry where conflict inputs are projected to different eyes, the pupillary responses can directly reflect the effectiveness of the differently suppressed stimulus. They found that the pupil constricted more to the sun pictures than to the scrambled controls only when the images were consciously perceived, not when they were suppressed. The results seem to suggest that conscious awareness of the pictures is necessary for the pupillary response to the interpreted brightness of the picture. However, an alternative possibility could be a failure of the brightness interpretation under the CFS condition, as the processing of high-level information is relatively shallow in interocular suppression (Gayet, Van der Stigchel, and Paffen, 2014; E. Yang, Brascamp, Kang, & Blake, 2014). Thus, the issue of whether conscious awareness of visual stimuli is necessary to trigger a proper pupillary response remains unsolved.

To better address this issue, we should find a suitable visual property that has already been proven to be processed under interocular suppression. Facial information, for example, discloses its special role in this regard. Behaviorally, Jiang et al. (2007) presented an upright face or inverted face under CFS while gradually increasing the faces’ contrast and asked observers to detect the face from CFS as soon as possible, the so-called breaking CFS procedure (b-CFS hereafter, Stein, Hebart, & Sterzer, 2011). They found that upright faces were detected faster than inverted ones. Interestingly, this inversion effect is much stronger for faces than for other familiar objects (Stein, Sterzer, & Peelen, 2012; Zhou, Zhang, Liu, Yang, & Qua, 2010), implying that it is not merely due to familiarity with orientation. Consistent with the behavioral results, neural image studies have shown that invisible faces, like visible ones, remain to trigger activation of the fusiform face area (FFA) (Jiang & He, 2006; Parallelly, Moutoussis & Zeki, 2002) and that decoding neural activation of the FFA can predict whether an invisible face is presented under the suppressed state or not (Sterzer, Haynes, Rees, 2008). Moreover, using magnetoencephalography (MEG), Sterzer, Jalkanen, and Rees (2009) found that an invisible face also induces a reliable M170 component, a face-related neural signal, like a visible face does, indicating neurological evidence of the particular role of the suppressed face. These convergent findings demonstrate a reliable and specific mechanism of facial information processing under interocular suppression, namely when the face is invisible.

In this study, we investigated pupillary responses to faces under different conscious awareness conditions in four experiments. Preconscious and unconscious/conscious statuses of faces were tested with the b-CFS procedure in Experiment 1 and 2 and with the fixed-duration CFS procedure in Experiment 3 and 4, respectively. To present the face and masks dichoptically under CFS, we used an infrared emitter to synchronize the left/right liquid crystal layer of shutter glasses with odd/even frames of a 120-Hz display. With this setting, a camera-based eye tracker can continuously record participants’ pupil sizes. In the b-CFS procedure in Experiment 1 and 2, participants had to detect the face as soon as it was released from interocular suppression. The stimulus processed under may go through a preconscious processing since this b-CFS procedure involves a transitory period in which the interocularly suppressed face gradually gains access to awareness (Gayet et al., 2014). Results showed that an upright face induced stronger pupil constriction than an inverted counterpart before the detection was made, suggesting that pupillary responses reflect face processing before the face is consciously perceived. In Experiment 3 and 4, we presented a face stimulus under CFS with a fixed duration and used both subjective reports and objective measurements to assess whether participants were aware of the face or not. We regarded the face as being *consciously* processed when participants reported subjectively high visibility/confidence and performed with high accuracy in the objective measurement, whereas we regarded the face as being *unconsciously* processed when participants reported subjectively low visibility/confidence and showed chance-level performance in the objective measurement. The results showed that stronger pupil constriction for the upright face than for the inverted face was observed when the face was consciously processed but not when it was unconsciously processed. Taken together, the overall results suggest that pupillary responses could be a precursor of the processing of faces in the suppression phase, i.e., preconscious processing; nevertheless, they cannot reflect the unconscious processing of faces.

## Results

### Pupillary reflection of preconscious processing of facial information (Experiment 1 and 2)

In Experiment 1, we tested how pupils respond to an upright face compared with the inverted counterpart face when the face gradually gains access to consciousness. To address this issue, we presented the face in one eye, which was interocularly suppressed by dynamic masks presented in the other eye (i.e., CFS), and gradually increased the face contrast (Fig. 1). Participants were asked to press the space key as soon as they had detected any part of the face (i.e., b-CFS) or hold their response if they did not see the face. To assess whether the participants pressed the key even when they did not see the face, we included catch trials in which no face was presented. The behavioral and pupillary responses were recorded at the same time.

**Figure 1.**
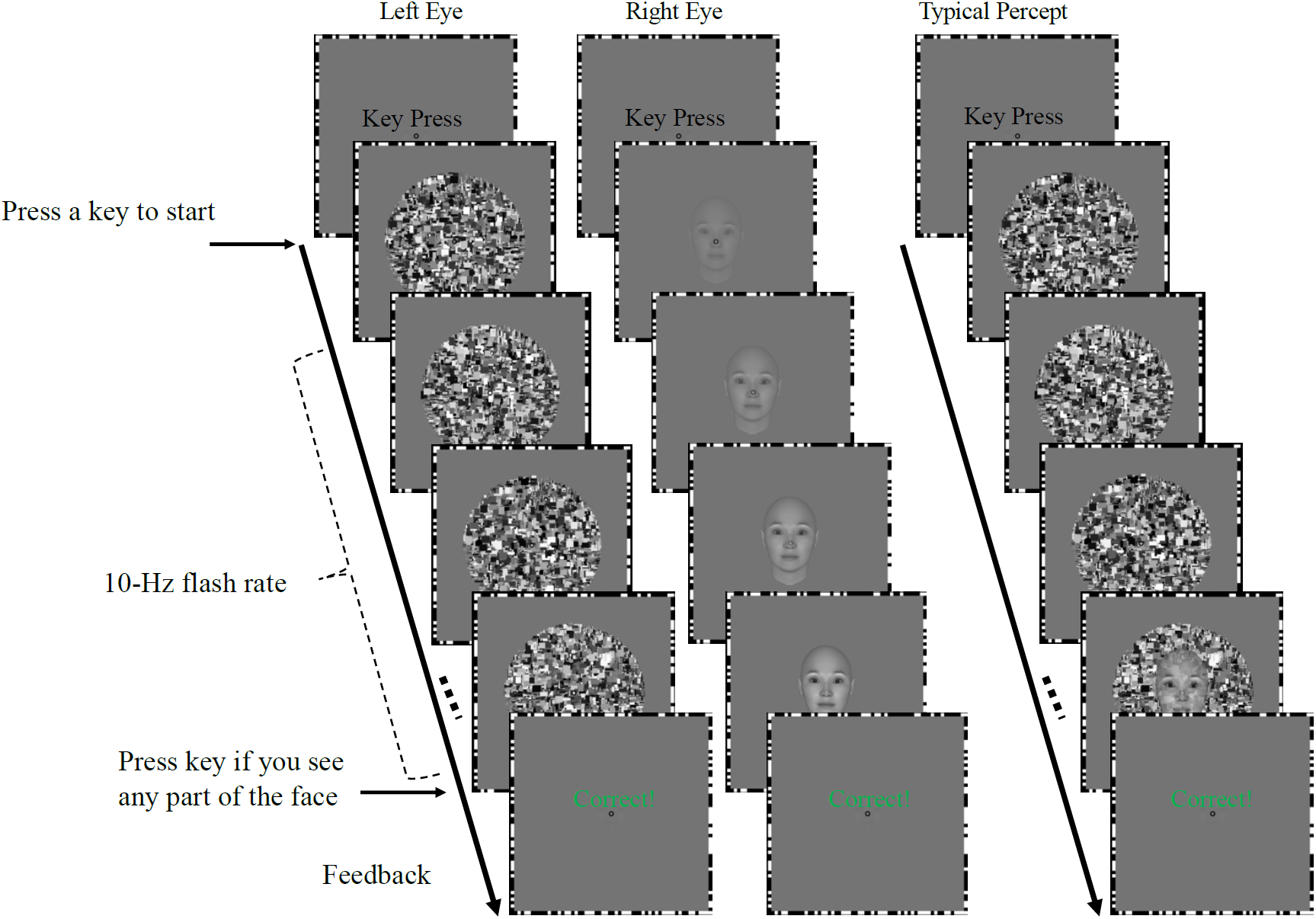
Procedure in Experiment 1. A key press started the trial. A face was presented to one eye, which was interocularly suppressed by flashing masks presented to the other eye. Participants were asked to press a key as soon as they detected any part of the face. After that, a feedback signal was presented. Reaction times and the pupillary responses were recorded.

On average, the accuracy was high (mean = 94.70%, SE = 2.11%), indicating that participants followed the instruction to press the key only when they detected the face. The upright faces (mean = 95.12 %, SE =1.89%) showed significantly higher accuracy (two-tailed, paired sample t-test; *t*(10) = 2.42, *p* = .0363, *Cohen’s d* = 0.765) than the inverted faces (mean = 92.59 %, SE =2.74%). Consistently, the upright faces (*M* = 1576 ms, *SE* = 143 ms) were detected faster (*t*(10) = 5.01, *p* = .0005, *Cohen’s d* = 1.595) than the inverted counterparts (*M* = 1811 ms; *SE* = 152 ms) (Figure 2, left panel). The result replicates the previously reported face inversion effect in the b-CFS procedure (Gray, Adams, Hedger, Newton, & Garner, 2013; Jiang et al., 2007; E. Yang Zald, & Blake, 2007; Y.-H. Yang & Yeh, 2018). Only trials with correct behavioral responses were included for further pupillary analysis. Pupil sizes as a function of time are shown in Fig. 2 (right panel) after stimulus onset, where the pupil size is expressed relatively to the mean value of the pre-stimulus period (−500 to 0 ms). The upright faces induced stronger pupillary constriction than the inverted face or no-face conditions. From a cluster-based permutation test (α < .05), we found significantly different clusters in the time window from 1148 to 1666 ms for the comparison between the upright and inverted faces and from 1059 to 2000 ms for the comparison between upright and no-face conditions. The result cannot be simply explained by low-level features such as contrast or luminance changes because the upright faces and inverted faces were the same faces but with different orientations. The finding that the onset of pupillary responses was faster than behavioral ones suggests that pupil sizes can reflect how participants processed the face before they became aware of facial stimuli under interocular suppression.

**Figure 2.**
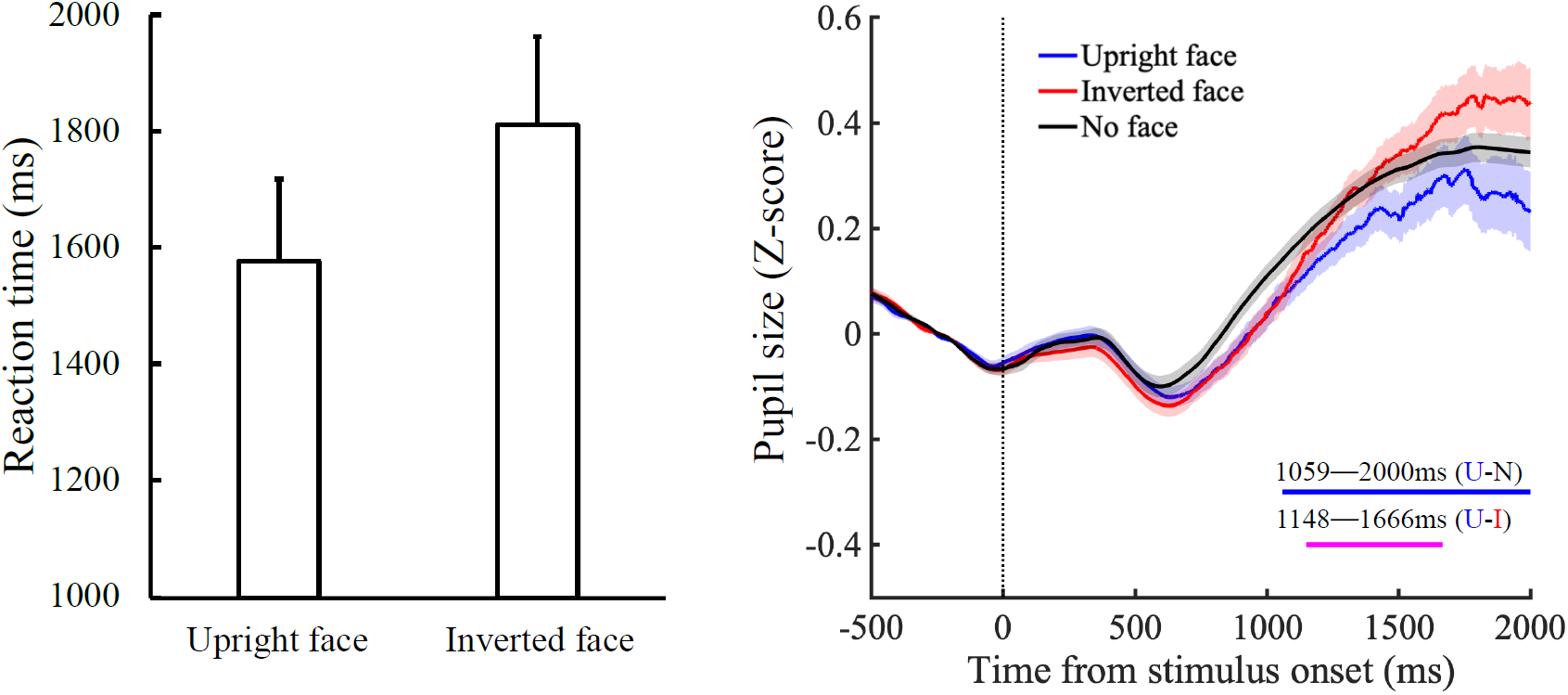
Results of Experiment 1. As shown in the left panel, upright faces had faster breaking-CFS time than inverted ones. As shown in the right panel, pupil sizes [corrected relative to the baseline (−500 to 0 ms)] indicate that upright faces induced a stronger pupil constriction response than inverted faces or the no-face condition. The horizontal magenta (upright face vs. inverted face) and blue lines (upright face vs. no face) represent significant clusters (α < .05), and the vertical dotted line represents stimulus onset.

We were concerned about the possibility that the relatively slow behavioral response found in Experiment 1 was attributable to a low arousal level and/or a bias towards conservative decisions by the participants: To cope with the no-face condition, in particular, a reasonable strategy for the participants would be to hold their responses for a relatively long period of time. Therefore, we designed Experiment 2 in which, as a measurement ensuring that participants indeed had detected any part of the face from CFS and then pressed the key, we used a face localization task instead of including the no-face condition. The procedure was similar to that in Experiment 1, but the face was presented to either left or right visual fields in every trial. Participants were asked to detect the face location under interocular suppression as soon and as accurately as possible. The accuracy of face localization showed no significant difference (two-tailed, paired sample t-test; *t*(11) = 1.590, *p* = .0140, *Cohen’s d* = 0.482) between the upright faces *(M* = 98.02%, *SE* = 0.84%) and the inverted faces *(M* = 97.17%, *SE* = 0.94%). Consistent with Experiment 1, we also found a clear face inversion effect in the reaction time. That is, upright faces *(M = 1316 ms, SE = 68 ms)* were detected significantly faster *(t*(11) = 3.92, *p* = .0024, *Cohen’s d* = 1.183) than inverted faces *(M = 1394 ms, SE = 72 ms)* (Fig. 3, left panel). Pupil size as a function of time is shown in the right panel of Fig. 3. Parallel to the behavior result, we found that upright faces induced stronger pupil constriction than inverted faces with a significant cluster after the onset of face presentation around 422–840 ms. We noted that the RTs and pupillary responses were generally faster in Experiment 2 than in Experiment 1. This may reflect the participants’ higher arousal and/or less conservative decisions as expected from the exclusion of the no-face condition (see beginning of this paragraph). On top of this finding, the convergent evidence from both b-CFS experiments shows that the pupil responded more strongly to upright faces than to inverted ones when the interocularly suppressed face gradually gained access to consciousness, which implies that pupillary responses could be an index of the preconscious processing of faces in the suppression phase.

**Figure 3.**
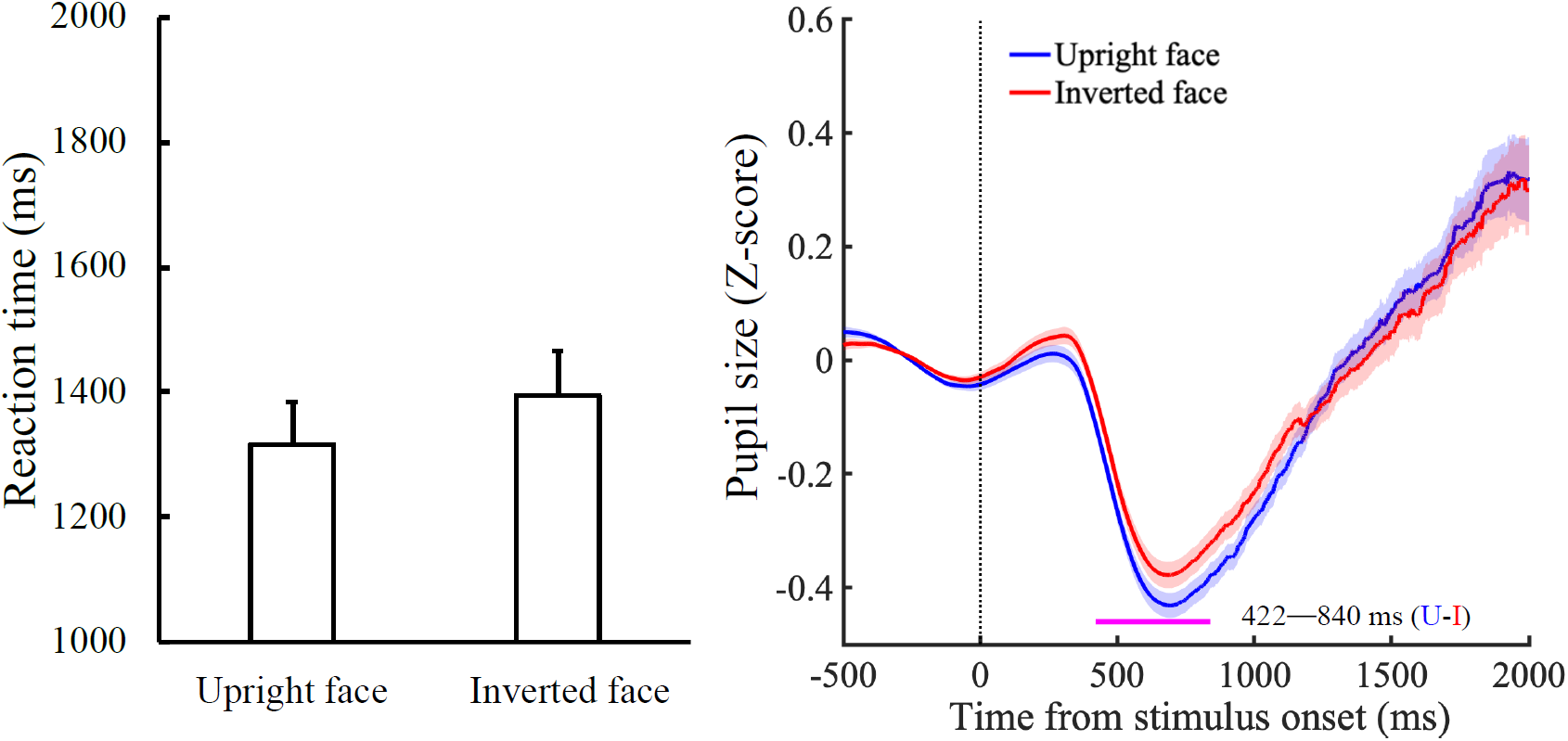
Results of Experiment 2. As shown in the left panel, upright faces had faster breaking-CFS time than inverted faces. As shown in the right panel, pupil sizes [corrected relative to the baseline (−500 to 0 ms)] indicate that upright faces induced stronger pupil constriction response than inverted faces. The horizontal magenta line represents significant clusters (α < .05) and the vertical dotted line represents stimulus onset.

### Pupillary reflection of conscious/unconscious processing of facial information (Experiment 3 and 4)

Experiment 1 and 2 indicate that pupil size reflected face orientation well before the participant became aware of the face. Does this mean that a pupil-related component as a part of the face-detection processing chain starts operating before the consciousness-related component, or that pupil- and consciousness-related mechanisms are effectively independent, i.e., the activation of the pupil-related mechanism does not have to be accompanied by the (eventual) activation of the consciousness-related mechanism? Experiment 3 addressed this issue by testing whether pupillary constriction for upright faces still occurs when the conscious awareness of faces is *fully* suppressed under CFS. We presented faces in the left or right visual field of the nondominant eye, and their contrast was continuously weakened for two seconds. In addition, to increase the motivation of participants, we added filler trials where the face contrast was high (Rothkirch, Stein, Sekutowicz, & Sterzer, 2012). To assess the conscious awareness of the face presentation, participants were asked to indicate the face location (left or right) and the confidence in their response (low, middle or high; see Materials and methods for details). By sorting the data by performance in the face localization task and confidence rating with different criteria, we regarded the conscious and unconscious processing of face information in the filler and CFS condition, respectively. In the filler condition, participants reported generally high confidence in their responses (proportion of confidence ratings: low, 9.8% ±19.7%; middle, 12.7% ±31.1%; high, 73.0% ±40.4%). In contrast, in the CFS condition, participants reported generally low confidence in their responses in most of the trials (proportion of total trials: low, 62.8% ±29.0%; middle, 12.7% ±15.6%; high, 23.5% ±28.3%).

For the low-confidence-rating trials of the CFS condition, accuracy for the face localization task showed performance significantly higher than chance level for both the upright (M = 59.03%, SD = 9.28%, t(19) = 4.35, p < .0001, *Cohen’s d* = 0.97) and inverted (M = 59.73%, SD = 8.13%, t(19) = 5.35, p < .0001, *Cohen’s d* = 1.2) faces, indicating that some participants were partially aware of the face location. Therefore, we used a binomial test for each participant and removed twelve participants who showed higher than chance performance (χ^*2*^*s*> 4.04, *p*s < .044). After the data had been trimmed, the accuracy for face localization task did not differ from chance level for both the upright (M = 50.84%, SD = 3.40%, t(7) = 0.70, p = .50, *Cohen’s d* = 0.24) and inverted faces (M = 53.12%, SD = 4.79%, t(7) = 1.84, p = .11, *Cohen’s d* = 0.65), suggesting the rest of the participants were fully unaware of face location.

The left panel of Fig. 4 shows the pupillary responses in the filler condition, in which only trials with “high” confidence ratings and correct responses in the face localization task were included to represent conscious processing. The upright faces showed stronger pupillary constriction than the inverted ones with a significant cluster from 1071 to 1582 ms. This result replicates previous findings that upright faces induce stronger pupillary constriction than inverted ones when participants are fully aware of the faces (Conway et al., 2008). In contrast, in the CFS condition (Fig. 4, right panel; only trials with “low” confidence ratings and for participants who’s face localization performance was at chance level were included), we did not find a significant cluster for comparison between upright and inverted face. This null result would suggest that pupillary responses cannot reflect the unconscious processing of faces.

**Figure 4.**
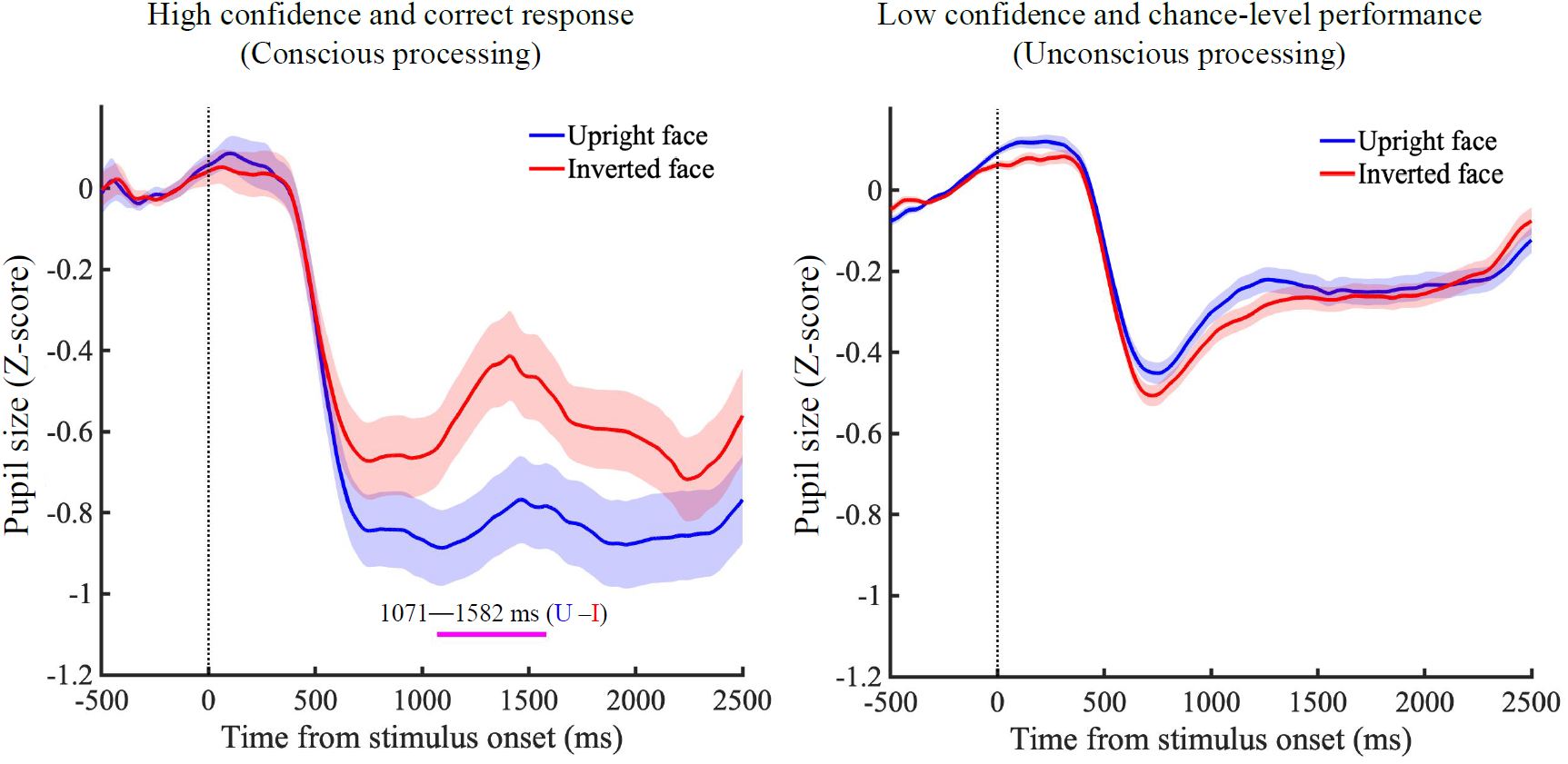
Pupillary responses to visual stimuli [corrected relatively to the baseline (−500 to 0 ms)] in Experiment 3. In the conscious trials (left panel), upright faces induced stronger pupil constriction than inverted faces for the trials with high confidence and correct responses in the filler condition. In the unconscious trials (right panel), there was no significant difference for participants who showed chance-level performance for the trials with low confidence in the CFS condition. The horizontal magenta line represents significant clusters (α < .05), and the vertical dotted line represents stimulus onset.

In this experiment, we tested whether upright and inverted faces are processed differently and failed to find evidence when the conscious awareness of faces is fully suppressed. One possibility is that the pupil-related mechanism was sensitive to faces but could not discriminate the face orientations in the absence of consciousness. In Experiment 4, we addressed this issue by including scrambled faces as a control condition, and tested whether the pupillary responses could reflect the difference between faces (i.e., upright or inverted faces) versus non-faces (scrambled faces) conditions.

In Experiment 4, besides upright faces and inverted faces, scrambled faces were included to serve as the control condition (see Fig. 5). Additionally, to better suppress the conscious awareness of faces, their edges were spatially smoothed so that they merged with the background to avoid a sharp boundary, and the face contrast was adjusted every trial based on the mean accuracy of a face orientation judgment task to make the performance chance level. As in Experiment 3, the filler condition was included to maintain the motivation of participants. Participants were asked to report their subjective opinion on the visibility of the faces (visible or invisible) and to judge face orientation (upright or inverted; see Materials and methods for details). Similar to Experiment 3, by sorting the data by performance in the face orientation task and subjective rating, we assumed that we could access the conscious and unconscious processing of face information in the filler and CFS condition, respectively.

**Figure 5.**
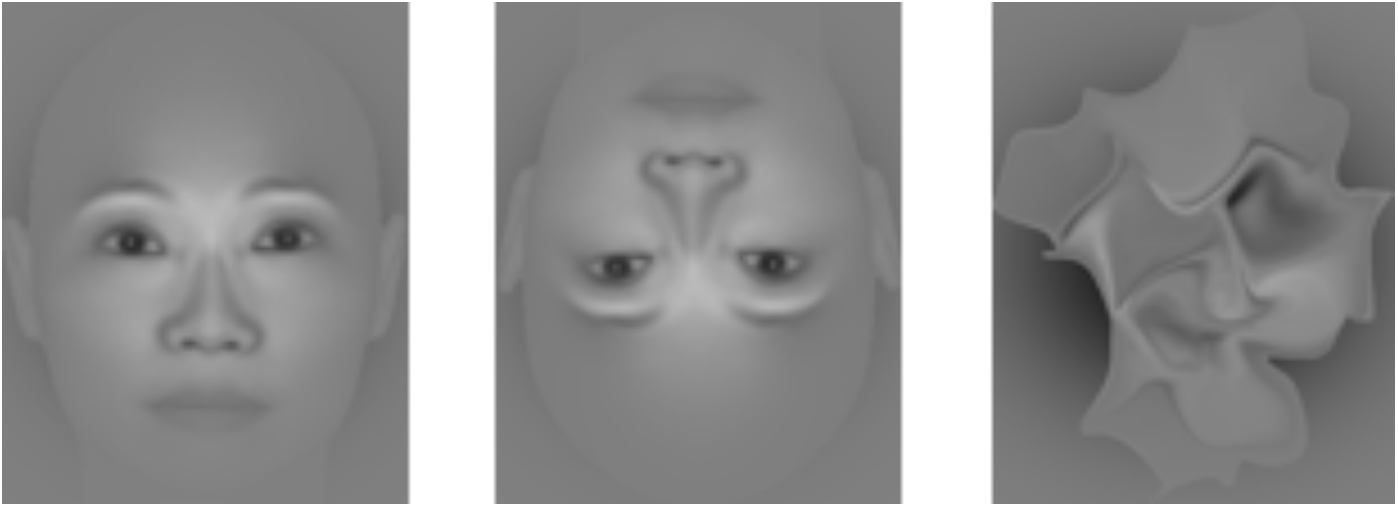
Visual stimuli used in Experiment 4. From left to right: upright, inverted, and scrambled faces.

In the filler condition, participants reported high visibility in most trials (proportion of visibility: upright, 96.88% ±10.49%; inverted, 89.06% ±18.93%; scrambled, 68.75% ±41.67%). In contrast, in the CFS condition, participants reported low visibility in most trials (proportion of visibility: upright, 4.40% ±9.69%; inverted, 3.65% ±8.91%; scrambled, 1.56% ±6.88%). The mean accuracy of face orientation judgment was not different from chance level in the CFS condition (t(15) = 0.569, p = .578, M = 50.50%, SD = 4.23%). Two participants who exhibited higher than chance performance (χ^*2*^> 5.16, p < .023, M = 58.48%, SD = 0.94%) were excluded from further analyses. For the pupillary response in the filler condition (Fig. 6, left panel), only “visible” trials and correct responses in the face orientation task were included to represent conscious processing. The result also shows that upright faces induced stronger pupillary constriction than inverted ones at a significant cluster from 693 to 2000 ms, which is consistent with previous findings (Conway et al., 2008) and the conscious trials in our Experiment 3. Unexpectedly, there was no difference between upright and scrambled faces, and pupillary constriction was stronger for scrambled faces than for inverted ones at a significant cluster from 1033 to 1802ms. This result may be related to the participants’ tendency to report a scrambled face as an upright one (M = 66.6%, SD= 23.8%) more frequently than as an inverted one (M= 33.3%, SD = 23.8%), and thus pupillary responses may reflect this perception of face orientations.

**Figure 6.**
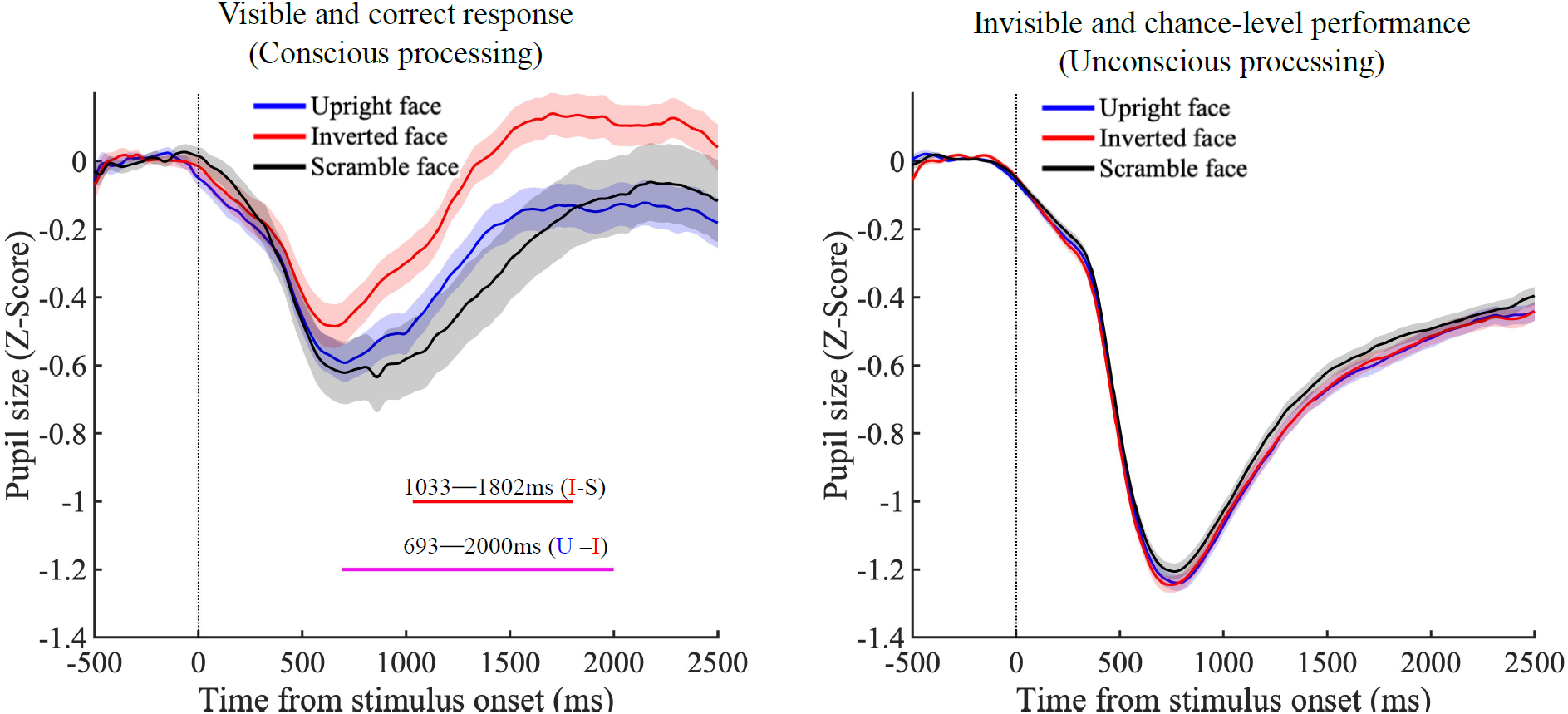
Pupillary responses to visual stimuli [corrected relatively to the baseline (−500 to 0 ms)] in Experiment 4. For the conscious trials (left panel), both the upright and scrambled faces induced a stronger pupil constriction response than the inverted faces for the visible and correct response trials in the filler condition. For the unconscious trials (right panel), there is no significant difference among upright, inverted, and scrambled faces for the invisible trials and participants who showed chance-level performance in the CFS condition. The horizontal magenta (upright face vs. inverted face) and red lines (scrambled face vs. inverted face) represent significant clusters (α < .05), and the vertical dotted line represents stimulus onset.

On the other hand, there was no significant differences in the pupillary responses among the face conditions in the CFS condition (Fig. 6, right panel). It should be noted that the CFS condition here represents trials with “invisible” responses only and participants whose face-orientation discrimination performance was at chance level. This null result implies that pupillary responses were neither sensitive to face orientation (i.e., upright faces versus inverted faces) nor face information (i.e., upright/inverted faces versus scrambled faces) when the consciousness to faces was fully suppressed under interocular suppression.

## Discussion

In the current study, we conducted four experiments to test pupillary responses to preconscious and unconscious facial information processing under interocular suppression. Experiment 1 and 2, which used the breaking-CFS paradigm (Jiang et al., 2007), indicated that observers detected the upright faces faster than their inverted counterparts. Correspondingly, the upright faces induced stronger pupillary constriction than the inverted faces. This results are in line with a previous study where participants were fully aware of the face (Conway et al., 2008). Our findings advance the finding by demonstrating that the effect occurs before the facial information reaches awareness. On the contrary, when the faces were entirely suppressed under interocular suppression (Experiment 3 and 4), the pupillary responses did not show any sensitivity to upright versus inverted faces.

Previous studies have shown that the pupillary responses are related to the conscious representation of perceptual changes in bistable stimuli (Einhäuser, Stout, Koch, & Carter, 2008; Hupé; Lamirel, & Lorenceau, 2009; Lamirel. Hupé, & Lorenceau, 2010). For example, Einhäuser et al. (2008) found that the pupil dilates slightly earlier than observers’ timing of button presses to indicate perceptual switches when viewing a Necker cube. The results of Experiment 1 and 2 add to their finding in that pupillary changes reflect not only the timing of the perceptual selection but also the *content* of the visual object entering consciousness. That is, the pupil constricts more when upright faces are being accessed to awareness compared to inverted faces, supporting the pupillary responses to the preconscious processing of face information. In this study, we found that distinct pupillary responses between the unconscious versus preconscious processing of face information. On the one hand, we did not find pupillary responses when face awareness was fully suppressed (Experiment 3 and 4). According to the model of pupillary responses proposed by Barbur (2004), the EW nucleus receives inhibition projections from the cortical areas, and the presentation of a visual stimulus decreases the inhibition projection and induces pupil constriction. If the pupillary response to faces is underlain by this sensory cortex pathway, regardless of whether the face is visible or invisible, we should observe similar pupillary responses in the conscious and unconscious conditions. Instead, our findings suggest that when the face’s conscious awareness is entirely blocked by interocular suppression, the projections from cortical areas seem to attenuate the modulation of pupillary constriction. This implies that not only the sensory cortex but also the prefrontal cortex is involved in the consciousness-related mechanism to induce pupillary responses (see more discussion about the locus-coeruleus norepinephrine (LC-NE) system below). On the other hand, we found pupillary responses when the suppressed face breaking through interocular suppression. It should be noted that the preconscious processing studied here was different from the unconscious processing in that the suppressed face reached conscious awareness eventually. In the b-CFS procedure, upright faces were detected more quickly than inverted faces, suggesting efficient processing of upright faces even during interocular suppression (Jiang et al., 2007; Stein et al., 2012; Zhou et al., 2010). This face inversion effect is well-documented in the literature for cases in which people are fully aware of the face (Civile, McLaren, & McLaren, 2014; Yin, 1969). Consistently, our results show that the upright face induced stronger pupil constriction than the inverted face preceding the conscious awareness of face presence. This implies that even though the pupil-related component in the face-detection processing acts faster than eventual conscious percepts, the activation of the pupil-related component must be accompanied by the activation of the consciousness-related mechanism during this preconscious processing. Moreover, when the consciousness-related mechanism is triggered, the pupillometric responses could serve as an objective measurement to assess both preconscious and conscious processing.

What is the possible physiological mechanism linking consciousness modulation and the pupillary responses? A candidate is the LC-NE system (Aston-Jones & Cohen, 2005; Bouret & Sara, 2005), which not only projects an excitatory signal to the superior cervical ganglion to control pupil dilatation via the iris dilator muscle but also projects an inhibitory signal to the EW to inhibit pupil constriction. Critically, the LC-NE system also receives inputs from subcortical regions such as the superior colliculus (Wang & Munoz 2015) as well as higher level cortical inputs from the anterior cingulate cortex (ACC), prefrontal cortex, etc. (Aston-Jones & Cohen, 2005; Josh, Li, Kalwani, Gold, 2016). As global recurrent from the prefrontal cortex may play an essential role in conscious experience (Dehaene et al., 2006), this area’s activation may modulate the LC-NE system and thus influence pupillary responses to face information.

Interestingly, while the present study showed no *pupillary* responses to facial information under interocular suppression, a recent study found that threat-related facial expressions can guide *oculomotor* responses in the absence of consciousness (Vetter, Badde, Phelps & Carrasco, 2019). Their research presented fearful, angry, and neutral faces under CFS and measured gaze direction directly. They found that gaze direction avoided an angry face (direct threat) but approached a fearful face (indirect threat) under full suppression. They thus suggested that affective facial expression might go through the subcortical retinocollicular pathway and superior colliculus to guide gaze direction in response to threat-related information. It is premature to conclude that the negative and positive evidence for the contribution of consciousness to face-related responses by the present and the Vetter et al.’s studies, respectively, reflects the existence of separate pathways for pupillary and oculomotor responses. The face stimuli adopted in the current study included only neutral faces and thus may not have triggered subcortical processes as emotional, affective faces would do (Jiang & He, 2006).

In summary, the current study sheds light on the mechanism of pupillary responses and conscious modulation. On the one hand, we found that pupillary responses required conscious representation of visual content, which is in line with previous findings showing that they are sensitive to conscious awareness (Fahle et al., 2011; Naber et al., 2011; Schütz et al., 2018; Sperandio et al., 2018). Critically, we also found that pupillary responses reflect different processing between upright and inverted faces before the face is consciously perceived. This finding has theoretical and methodological implications for our understanding of preconscious processing. Pupillary responses may provide information about dynamic aspects of conscious awareness, in addition to the knowledge obtained from typical behavioral reaction times in the b-CFS paradigm.

## Materials and methods

### Participants

Fifty-nine observers who were naïve about the purpose of the experiment took part in this study (Experiment 1 (n=11), Experiment 2 (n=12), Experiment 3 (n=20), and Experiment 4 (n=16)). Each participant had normal or corrected-to-normal visual acuity. The experiment was approved by the Ethics Committee of the Nippon Telegraph and Telephone Corporation, Japan (H30-011). All participants gave informed written consent before the experiment.

### Visual stimuli and apparatus

In all the experiments, visual stimuli were rendered in Matlab R2018a (MathWorks, Natick, USA) using Psychophysics Toolbox Version 3 and were presented on a 24-inch Asus VG248QE (1,920×1,080 resolution at 120-Hz refresh rate). Active shutter glasses (Nvidia 3D vision 2) were synchronized with the refresh rate of the monitor by an infrared emitter so that odd and even frames presented the stimuli dichoptically.

An outer frame (28° × 28°) with black-and-white interleaved rectangles was presented on a gray background ([128 128 128] on RGB scale) to each eye for stable binocular fusion. A small empty circle (radius: 0.7°) in the center of the display served as a fixation point for both eyes. A Mondrian-like circle (radius: 10°) containing 3,000 rectangle patches (random grayscale: 0∼255; random size: 0.02° to 1.10°) was presented to one eye to serve as a suppressing mask, and a face target (7° × 7°) was presented to the other eye. The face stimuli were generated from FaceGen Modeller (Singular Inversions, Toronto, Canada) and included six posers (three female) with a neutral facial expression. To create inverted faces, we rotated the same set of upright faces 180°. With this manipulation, the upright and inverted faces had the same physical properties. Face stimuli were presented to the center of the visual field in Experiment 1 and 4. In Experiment 2 and 3, they were presented in either the left or right visual field with equal chance for the face localization task (see procedure), and the center-to-center distance between the face and fixation was 5°.

In the b-CFS tasks (Experiment 1 and 2), we increased the contrast of the face gradually from 0 to 100% for one second and kept it constant afterward to prevent an abrupt onset of the face from inducing a dominant percept at the beginning. To control the ocular dominance, we also counterbalanced the eyes to which the faces or masks were presented. In Experiment 3, to keep faces invisible during CFS, we presented them to the nondominant eye, and the contrast of the face was ramped up gradually from 0% to 20% within the first second and kept constant for another one second. Since more than half of the participants showed higher than chance performance in Experiment 3, we modified the visual properties of the stimuli to better suppress the face in Experiment 4. We used spatial smoothing to make the edges of faces blend in with the gray background. This was done to prevent sharp facial boundaries from being detected and thus better suppress the conscious awareness of faces. More specifically, we adjusted the contrast of the faces using an adaptive contrast procedure based on the mean accuracy of the face orientation task after every trial (Y.-H. Yang et al., 2017). On average, the contrast of faces was 50.14 % (SD= 9.16 %). Additionally, since the face was supposedly entirely suppressed in Experiment 3 and 4, we conducted filler trials where the face contrast was 100% starting from the beginning of the trials to increase the motivation of participants (Rothkirch et al., 2012).

### Experimental Design

All four experiments had three repeated sessions (one participant only finished two sessions due to the time limit in Experiment 1), but trial numbers and conditions within sessions were varied depended on the experimental designs. In Experiment 1, we used an within-subject design with a single factor of face presentation (upright, inverted, and no face). The no-face condition served as catch trials to assess whether the participants pressed the key even when they did not see a face. Each session contained 108 trials and 36 trials per condition. In Experiment 2, we used an within-subject design with face orientation (upright and inverted face). Unlike Experiment 1, the no-face (catch trials) condition was excluded to prevent potentially low arousal/ conservative decisions. Each session contained 108 trials and 54 trials per condition. In Experiment 3, we used a within-subject design with face orientation (upright and inverted face). Each session consisted of 132 trials: 60 CFS trials and six filler trials for upright and inverted face, respectively. In Experiment 4, we used a within-subject design with face presentation (upright, inverted, and scrambled face). As can be seen in Fig. 5, the scrambled faces were generated from the upright faces by diffeomorphic transformation (Stojanoski, & Cusack, 2014) to serve as the baseline. The orientation of scrambled faces was randomly assigned as upright (0°) and the inverted (180°) counterpart. Each session contained 120 trials: 36 CFS trials and four filler trials for each type of face (upright, inverted, and scrambled), respectively. In all experiments, the sequence of conditions was pseudorandomized within each session.

### Procedure

In all the experiments, after participants pressed the space key to start each trial, and a critical visual stimulus and a series of masks were presented dichoptically. In Experiment 1, participants were asked to press the space key as soon and as accurately as they detected any part of the face or to hold their response if they did not see the face. In Experiment 2, participants were asked to press the left-arrow key and right-arrow key as soon as possible once they had detected the face that was presented in either the left or right visual field, respectively. In both b-CFS tasks (i.e., Experiment 1 and 2), the trial was ended immediately after participants had pressed the key or after six seconds without a key press. After that, a feedback frame with either “correct” (green) or “wrong” (red) was presented (Fig. 1). In Experiment 3, a face (upright or inverted face) was either presented in the left or right visual field, in which the face was interocularly suppressed by a series of masks for two seconds. After the face presentation, participants were asked two questions: (1) Where did you think the stimulus was presented, left or right? They pressed the number “4” key and number “6” key in the numeric keypad to indicate the left and right responses, respectively. (2) How confident are you about your answer about the stimulus location? They pressed the key number “1”,” 2” and “3” key to indicate a low, middle and high confidence rating, respectively (Kanai, Walsh, & Tseng, 2010). Unlike previous experiments, no feedback was presented. In Experiment 4, a face (upright, inverted, or scrambled face) was presented in the center, where it was interocularly suppressed by a series of masks for two seconds. Afterward, participants were asked to answer two questions: (1) Did you see any parts of the face? (2) Was the face upright or inverted? The formal served as a subjective criterion of unawareness; the latter served as an objective measurement of unawareness. An adaptive contrast procedure was also included in this experiment to adjust contrast for face suppression; that is, after each trial, we calculated the mean accuracy from upright faces and inverted faces (the scrambled faces were not included). The contrast of faces was decreased by 2% if the mean accuracy was higher than 50% and increased by 2% if the mean accuracy was lower than 50%.

### Pupillary *Recording and Statistical Analysis*

Throughout all the experiments, pupillary responses were recorded monocularly using an eye-tracking system (Eyelink 1000 Desktop Mount, SR Research, Mississauga, Canada) with a sampling rate of 1000 Hz. Five-point calibration and validation were conducted before each session. Blinks were detected by the default algorithm of the EyeLink system and treated as missing data. Data during a period of 50 ms before and after a blink were also treated as missing data to prevent artifacts from the closing and reopening of the eyelid, respectively.

Pupil sizes were time-locked to the onset of stimuli. To reconstruct pupil sizes, we conducted a cubic-spline interpolation for the missing data (Mathôt, 2013). After interpolation, pupil sizes were low-pass filtered with third-order Butterworth filters with a 10-Hz cutoff frequency to remove spikes from pupil data. We also excluded the full trial if the blink rate was higher than 20% of the trial duration. To conduct a baseline correction, we averaged pupil size during a period of 500 ms before the onset of each trial and subtracted the baseline pupil size from the pupil size data for each trial (Mathôt, Fabius, Van Heusden, & Van der Stigchel, 2018). For the statistical analysis, we adopted a cluster-based permutation test (Zuber, Stark, & Cook, 1965) with the FieldTrip toolbox (Oostenveld, Fries, Maris, & Schoffelen, 2010) to deal with the multiple comparisons problem as the comparison between two conditions relied on a large number of data points across time series.

## Notes

### Competing Interest Statement

The authors have declared no competing interest.

